# Diverse phenotypic and genetic responses to short-term selection in evolving *Escherichia coli* populations

**DOI:** 10.1101/027086

**Authors:** Marcus M. Dillon, Nicholas P. Rouillard, Brian Van Dam, Romain Gallet, Vaughn S. Cooper

**Affiliations:** Department of Molecular, Cellular, and Biomedical Sciences, University of New Hampshire, 105 Main Street, Durham, NH, 03824; INRA – UMR BGPI Cirad TA A-54/K Campus International de Baillarguet 34398 Montpellier Cedex 5 FRANCE; Department of Microbiology and Molecular Genetics, University of Pittsburgh School of Medicine, 425 Bridgeside Pt II, 450 Technology Dr, Pittsburgh, PA 15219

**Keywords:** beneficial mutations, pleiotropy, niche breadth, adaptive history, adaptation

## Abstract

Beneficial mutations fuel adaptation by altering phenotypes that enhance the fit of organisms to their environment. However, the phenotypic effects of mutations often depend on ecological context, making the distribution of effects across multiple environments essential to understanding the true nature of beneficial mutations. Studies that address both the genetic basis and ecological consequences of adaptive mutations remain rare. Here, we characterize the direct and pleiotropic fitness effects of a collection of 21 first-step beneficial mutants derived from naïve and adapted genotypes used in a long-term experimental evolution of *Escherichia coli*. Whole-genome sequencing was used to identify most beneficial mutations. In contrast to previous studies, we find diverse fitness effects of mutations selected in a simple environment and few cases of genetic parallelism. The pleiotropic effects of these mutations were predominantly positive but some mutants were highly antagonistic in alternative environments. Further, the fitness effects of mutations derived from the adapted genotypes were dramatically reduced in nearly all environments. These findings suggest that many beneficial variants are accessible from a single point on the fitness landscape, and the fixation of alternative beneficial mutations may have dramatic consequences for niche breadth reduction via metabolic erosion.

## INTRODUCTION

Adaptation proceeds through the accumulation of beneficial mutations that provide an advantage in the selective environment. The magnitude of the fitness effects provided by beneficial mutations typically declines as organisms adapt (Khan et al. 2011; Wiser et al. 2013; Kryazhimskiy et al. 2014), but how adaptation changes the shape of the distribution of the fitness benefits is less certain (Kassen and Bataillon 2006; Martin and Lenormand 2008; Good et al. 2012). In addition, the molecular targets of beneficial mutations appear to be relatively limited in the early steps of adaptation (Travisano et al. 1995b; Woods et al. 2006; Ostrowski et al. 2008), but may vary as evolution proceeds (Barrick et al. 2009). Both the direct (selected) and indirect (pleiotropic) effects of beneficial mutations are of central importance to countless biological processes, including adaptation (Cooper 2002; Leiby and Marx 2014), speciation (Cooper and Lenski 2000; Otto 2004), senescence (Holt 1996), antibiotic resistance (Kassen and Bataillon 2006; Bataillon et al. 2011), and the emergence of new pathogens (Kassen and Bataillon 2006). Therefore, characterizing the direct and pleiotropic effects of beneficial mutations as organisms adapt, along with their molecular targets, is fundamental to our understanding of adaptive evolution (Nahum et al. 2015).

While all aspects of the nature of beneficial mutations require further study, the environmental dependence of both the magnitude and sign of fitness effects has received especially little consideration, despite evidence that pleiotropy is abundant and possibly even universal (Cooper and Lenski 2000; Cooper et al. 2001; Rozen et al. 2002; Dudley et al. 2005; Ostrowski et al. 2005; Kassen and Bataillon 2006). Historical models of adaptation have often incorporated pleiotropy, but make several assumptions about the nature of pleiotropy that lack experimental support. Specifically, theoretical models often assume that pleiotropy is a largely antagonistic process, and that large effect mutations are more predisposed to antagonistic pleiotropy than small effect mutations (Fisher 1930; Lande 1983; Orr and Coyne 1992; Otto 2004). This framework has been invoked to explain the evolutionary success of small effect mutations, the variability of evolutionary trajectories, and why we see such immense diversity in nature despite the phenotypic parallelism observed in replicate laboratory populations (Travisano and Lenski 1996; Cooper et al. 2003; Ostrowski et al. 2008). However, too few data exist on the distribution of pleiotropic effects to assess the validity of these assumptions under a variety of population genetic environments (Good et al. 2012). As such, it is essential that we attain a more complete understanding of the nature of beneficial mutations and their pleiotropic effects.

The long-term evolution experiment (LTEE) with *Escherichia coli* provides an optimal system for studying the nature of beneficial mutations and their pleiotropic effects for several reasons. Firstly, the founding *E. coli* strain has a rapid replication rate and well-characterized genetics and metabolism (Cooper 2002; Feist et al. 2007; Jeong et al. 2009). Secondly, effective techniques for storage, measuring fitness, and manipulation of effective population size (N_E_) have already been established in this system (Lenski et al. 1991; Ostrowski et al. 2005; Gallet et al. 2012). Thirdly, the LTEE consists of a well-characterized evolutionary trajectory with rich genetic and phenotypic resources from previous studies, allowing for indirect manipulation of founding strains and a context for comparisons with previous studies. Lastly, of the few prior studies that have attempted to evaluate the direct and pleiotropic effects of individual beneficial mutations, several have been conducted in this system (Rozen et al. 2002; Ostrowski et al. 2005, 2008), allowing us to make direct comparisons between our findings and previous research.

Previous research in this system and others has yielded several insights into the direct and pleiotropic effects of beneficial mutations, but no single model consistently predicts how beneficial mutations in one environment cause pleiotropic effects in other environments (Travisano et al. 1995b; Ostrowski et al. 2005). Several genetic and phenotypic targets of adaptation have been identified in the LTEE with a high degree of molecular parallelism (Travisano et al. 1995b; Travisano and Lenski 1996; Woods et al. 2006; Ostrowski et al. 2008; Barrick et al. 2009). Furthermore, even when genetic parallelism is low, phenotypic parallelism appears to be high when replicate lineages are adapted to the same environment at high effective population size (N_E_) (Le Gac et al. 2013). Over the course of 20,000 generations of adaptation to glucose, all twelve replicate populations in the LTEE experienced metabolic decay of unused resources, particularly early in the experiment when the rate of adaptation was most rapid (Cooper and Lenski 2000). While these studies suggests that antagonistic pleiotropy plays a central role in the niche breadth reduction experienced by these populations, subsequent analysis of these lineages at 50,000 generations revealed that the extent of metabolic decay was much less pronounced than previously thought, and that long-term metabolic erosion is primarily driven by mutation accumulation (Cooper 2014; Leiby and Marx 2014).

A limited role of antagonistic pleiotropy is also supported by a collection of mutants captured following short-term selection in the LTEE environment, since most pleiotropic effects were positive and correlated positively with the direct effects of mutations (Ostrowski et al. 2005). However, some examples of antagonistic pleiotropy were also discovered, particularly in alternative environments whose method of catabolism and uptake were most different from glucose, and these antagonistic effects did not correlate with the magnitude of the direct effect (Ostrowski et al. 2005).

Sequencing of loci known to be under selection in this system revealed mutations in most isolates, suggesting that selection had acted on these few targets, but other unsequenced loci may have also been targets of selection (Ostrowski et al. 2008).

While a few comprehensive studies have now been conducted to characterize the pleiotropic effects of adaptation, they suffer from one of two primary shortcomings: a) they involve mostly large libraries of induced mutations (Remold and Lenski 2001; Bataillon et al. 2011; Hietpas et al. 2013), or b) they involve the study of combinations of mutations (often unknown) in each genetic background (Travisano et al. 1995a,b; Cooper and Lenski 2000; Jasmin and Zeyl 2013; Leiby and Marx 2014). There remains very little data on the direct and pleiotropic effects of individual, naturally arising beneficial mutations, especially where the genetic identity is also known.

Here, we used short-term selection to isolate twenty-one mutations associated with improved fitness in a glucose-limited environment and characterized their fitness effects on five alternative carbon substrates. Beyond adding to our limited understanding of the distribution of fitness effects of individual beneficial mutations (Ostrowski et al. 2005, 2008; Bataillon et al. 2011), our study differs from previous work in three fundamental ways. First, we adapted our populations at lower effective population sizes (N_E_) than the original LTEE and isolated mutations from both the winning and losing fraction of evolving populations. These modifications were intended to increase both the variance among sampled beneficial mutations and the probability of picking mutants of lesser benefit. Second, we isolated beneficial mutations from both the LTEE ancestor and an evolved clone with an approximately 30% fitness increase relative to the ancestor. This design allowed us to study of the role of adaptive history on the effects of subsequent beneficial mutations. Lastly, we used whole-genome sequencing to identify the molecular basis of the beneficial mutations that led to these phenotypic changes and to associate specific mutations with their fitness effects. Overall, this project sought to broaden sampling of beneficial mutations, which are typically skewed towards those of large-effect (Gerrish and Lenski 1998; Orr 2003; Ostrowski et al. 2005, 2008; de Visser and Rozen 2006). We identified several novel molecular and phenotypic outcomes of adaptation to a simple glucose-limited laboratory environment, and present a more comprehensive perspective of the spectra of beneficial mutations and their pleiotropic effects.

## MATERIALS AND METHODS

## Bacterial Strains and Culture Conditions

Two strains were used to initiate short-term selection experiments from naïve and glucose-adapted genotypes. The naïve genotype was the ancestral *E. coli* B ancestor used in the LTEE, REL606 (Lenski et al. 1991), which has been cured of all active plasmids and bacteriophage. The glucose-adapted genotype was isolated after 2,000 generations of selection in the LTEE. This strain, named REL1206, is ∼30% more fit than *E. coli* REL606 as a result of at least five adaptive mutations (*rbsR*, *topA*, *spoT*, *glmUS*, and *pykF*) (Lenski et al. 1991; Barrick et al. 2009; Khan et al. 2011; Flynn et al. 2013).

We used flow cytometry to rapidly detect subtle shifts in population structure due to an arising beneficial mutation in our selection experiments, requiring that we chromosomally mark each of our ancestral strains with both cyan (CFP) and yellow (YFP) fluorescent proteins, as described in (Gallet et al. 2012). Briefly, CFP and YFP genes containing the P_A1_ promoter and a kanamycin resistance cassette were amplified using primers with 50 bases homologous to the *E. coli rhaA* locus at their 5’ ends.

These products were then independently transformed into both REL606 and REL1206, both bearing the pKD46 plasmid, leading to the insertion of YFP-Kan and CFP-Kan into their *rhaA* locus (Datsenko and Wanner 2000). After the removal of the thermosensitive pKD46 plasmid, the pCP20 plasmid was electroporated into each of the four resultant strains and heat shocked, leading to the removal of the Kan^R^ cassettes and four novel fluorescently marked strains, hereafter referred to as REL606-CFP, REL606-YFP, REL1206-CFP, and REL1206-YFP (Datsenko and Wanner 2000; Gallet et al. 2012).

All selection experiments and competitions were carried out in Davis Minimal Broth supplemented with 25 mg/L of the appropriate carbon substrate (DM25) (Lenski et al. 1991; Ostrowski et al. 2005). Mutants recovered from freezer stock were preconditioned in LB for 24h and then diluted 1:10,000 into DM25 for an additional 24h of growth prior to initiating selection or competition experiments. In contrast to the LTEE, which was carried out using 50ml glass flasks containing 10 ml of DM25 medium on a shaking incubator at 120 rpm, we conducted selections and competitions in 18x150mm glass capped tubes containing 5 ml of DM25 medium maintained in a roller drum at 30 rpm. In addition, during experimental evolution, all serial transfers were done using 10,000-fold dilutions to reduce N_E_.

### Mutant Collection and Isolation

The rate and likelihood of capturing any given beneficial mutation in these short-term selection experiments depends on the beneficial mutation rate (μ_b_), the effective population size (N_E_), and the number of generations until detection. Both theory and experiments support the notion that evolving populations at smaller N_E_, at least down to the point at which μN = 0.01, will effectively sample a more variable set of beneficial mutations and allow for the capture of more mutations of smaller effect (Gerrish and Lenski 1998; Wahl and Gerrish 2001; de Visser and Rozen 2005). Thus, in addition to using a smaller culture volume during our short-term selection experiments, we reduced the transfer bottleneck and hence N_E_ by 100-fold from the LTEE conditions.

Selection experiments were conducted as follows. Following independent acclimation of each ancestral culture, twelve co-cultures of oppositely marked REL606-CFP and REL606-YFP, and twelve co-cultures of oppositely marked REL1206-CFP and REL1206-YFP were initiated via 10,000-fold dilution. As a result of the small fitness differences in the oppositely marked strains (SI Text), the twelve REL606 populations were initiated with a 3:1 ratio of CFP to YFP, and the twelve REL1206 populations were initiated with a 9:1 ratio of CFP to YFP. All twenty-four populations were then passaged by daily 10,000-fold dilutions into fresh DM25-glucose media, measuring the relative frequency of the two markers every three days using a Millipore-8HT flow cytometer, until a marker divergence was observed or populations underwent 600 generations of selection (Figure S2; Figure S3).

Once these selection experiments were completed, populations were streaked onto LB plates and incubated overnight at 37°C. A clone from both the winning and losing marker states was selected, grown overnight in LB at 37°C, and stored at -80°C in 8% DMSO. This resulted in the collection of a total of forty-eight putative beneficial mutants (one winner and one loser from each population). However, for REL1206, marker divergence only occurred in four of the twelve populations, reducing the probability that the isolates actually harbored beneficial mutations. The relative fitness of each of these forty-eight isolates was measured relative to the oppositely marked ancestor with two-fold replication, and if increased fitness was observed, the isolate was kept for further analysis. These fitness assays identified a total of twenty-four putative beneficial mutations, including sixteen derived from the REL606 background and eight derived from the REL1206 background.

### Direct and Indirect Fitness Assays

The relative fitness of each of the twenty-four mutants was measured in direct competition with the oppositely marked ancestral strain of the short-term selection experiments using DM25-glucose, DM25-N-acetyl-d-glucosamine (NAG), DM25-trehalose, DM25-galactose, DM25-melibiose, and DM25-maltose. We chose to vary carbon substrates rather than other elements of the environment because of the wealth of physiological knowledge about how these substrates are processed in *E. coli*, and evidence from previous studies showing that substrate transport is a central target of selection early in the LTEE (Travisano and Lenski 1996; Ostrowski et al. 2005). Resources were chosen to cover a broad range of potential mechanisms of outer and inner membrane transport mechanisms. Glucose, NAG, galactose, and melibiose all use the OmpF porin for outer membrane transport, while trehalose and maltose use the LamB porin. For inner membrane transport, glucose, NAG, and trehalose all utilize the phosphotransferase system (PTS), while galactose, melibiose, and maltose use non-PTS pathways (Travisano and Lenski 1996; Ostrowski et al. 2005).

Fitness assays were carried out with each of the 24 beneficial mutants being assayed in all six alternative environments in four independent experiments (144 fitness assays per experiment, and 144 x 4 = 576 assays in total). Both competitors were resurrected from -80°C and acclimated to the experimental environment as described above. Competitors were then inoculated at a 1:1 ratio via a 100-fold dilution into fresh media, and the relative concentrations of each competitor were measured using a Millipore-8HT flow cytometer (5000 events per sample). Each competition was then maintained at 37°C for 72 hrs, with a 100-fold dilution into fresh DM25 every 24 hrs. At the end of the third competition cycle, the relative concentrations of each competitor were again measured via flow cytometry, and fitness was calculated as described previously in this system using the ratio of the realized Malthusian parameters of the two competitors (Lenski et al. 1991; Ostrowski et al. 2005, 2008). In addition, raw cell counts at the beginning (N_i_) and the end (N_f_) of all competitions, along with relative growth rates (Δr), generations elapsed by the reference strain (G), and selection coefficients (s) for each competition are provided in Supplementary Dataset 1 to enable comparison with studies using different metrics of competitive fitness (Chevin 2011; Perfeito et al. 2014).

### DNA Sequencing and Analysis

Genomic DNA was extracted from each of the twenty-four putative beneficial mutations using the Wizard Genomic DNA Purification Kit (Promega Inc.). Following library preparation using a modified Illumina Nextera protocol (Baym et al. 2015), sequencing was performed using the rapid run, 151-bp, paired-end mode on an Illumina HiSeq2500 at the University of New Hampshire Hubbard Center for Genomic Studies. Raw reads were processed using Trimmomatic to remove Nextera PE adapter sequences and quality control was performed on all sequences with fastQC (Andrews 2010; Bolger et al. 2014).

Following processing of the raw sequencing reads, the four output fastQ files from Trimmomatic were passaged through the breseq pipeline to map processed reads from each isolate to the *E. coli* REL606 reference genome and identify mutations using default settings (Deatherage and Barrick 2014). An average of 97% of the reads from each isolate were mapped to the reference genome, resulting in an average coverage of ≈80x per isolate (Table S2). We first focused on high-confidence mutations for each isolate by extracting the raw calls from each isolate and comparing them to all other isolates with the same genetic background. If any mutation was called in more than 50% of the isolates derived from the same genetic background, it was considered a fixed progenitor mutation and was discarded, which is justified given the limited parallelism at the nucleotide level seen in previous studies, even when gene level parallelism is high (Ostrowski et al. 2008; Barrick and Lenski 2009). Next, we examined reports from Unassigned Coverage, New Junction Evidence, and Marginal Read Alignments for potential missed mutations in lower confidence regions or structural variants that are more difficult to identify. We searched these lists for evidence of any of the genes that have previously been identified as common targets of selection in this system, including *rbs*, *spoT*, *glmU*, *topA*, *pykF*, *nadR*, *pbp-radA*, and *hokB/sokB*, as well as mutations picked up in other lineages of the LTEE (Cooper et al. 2001, Cooper et al. 2003; Woods et al. 2006; Ostrowski et al. 2008; Barrick et al. 2009). We also kept marginal calls for manual validation if at least ten reads covered the site and there was more than 80% consensus for a novel nucleotide. Lastly, we manually validated all mutations using breseq’s graphical output to verify our final set of mutations, a strategy that has previously been demonstrated to reduce the likelihood of false positives (Tenaillon et al. 2012). Manual validation led to us to discard the only two putative mutations that had been identified as marginal, resulting in a final mutation collection comprised only of high-confidence mutation calls.

### Statistical Analysis

Following completion of all fitness assays and sequencing, only twenty-one (fifteen REL606 derived; six REL1206 derived) of the initial twenty-four mutants were used for the analyses presented below because the remaining three lacked any genetic or phenotypic evidence that they harbored a beneficial mutation. Specifically, the relative fitness of one REL606-derived genotype and two REL1206-derived genotypes was not significantly different from 1, independent of multiple comparisons, in any of the six environments tested (two-tailed t-test), and had no mutations identified in the whole-genome sequencing analysis pipeline described above.

All statistical analyses were performed in R Version 0.98.1091 using the Stats analysis package (R Development Core Team 2013). Independent two-tailed t-tests were used to test whether the average effect of each mutant in each environment differed significantly from 1 at a threshold p-value of 0.05. However, because there were 126 experimental conditions (21 mutants × 6 resources), we applied a Benjamini-Hochberg correction to all of our p-values to ensure that our false positive rate was below 5% (Benjamini and Hochberg 1995). Analyses of variance (ANOVAs) were performed on the direct effects of the beneficial mutations on glucose to test for effects of mutant fitness and block (date). We also performed ANOVAs on the entire array of pleiotropic effects of our beneficial mutations to test for effects of mutant fitness, resource, and mutant*resource interaction. Linear regressions were used to evaluate correlations between the direct effect of each mutation on glucose and their pleiotropic effects on each alternative resource. Shapiro-Wilkes tests were used to test for normality in the distribution of fitness effects for each environment. When the distribution was significantly non-normal, non-parametric Kruskal-Wallis tests were used to accompany ANOVAs, and non-parametric Spearman’s Rank correlations were conducted to accompany linear regressions.

### RESULTS

Experimental evolutions derived from the naïve REL606 genotype were evolved for between 130 to 300 generations, at which point sufficient deviation from the marker-generated trajectory was observed to warrant the inference that a beneficial mutation had arisen in at least one of the oppositely marked populations (Figure S2). Deviation from the marker-generated trajectory took substantially longer in the glucose-adapted REL1206 lineages (300-600 generations), and several populations did not diverge at all (Figure S3). When sufficient deviation was observed, a mutant was isolated from both the winning and losing marker portions of the population. For the REL1206 lineages that did not diverge from the marker-predicted trajectory after ≈540 generations, the experiments were stopped and putative mutants were isolated from both marker portions.

**Table 1:** Molecular basis and COG Cluster of all beneficial mutations isolated in this study from the naïve REL606 genotype and the glucose-adapted REL1206 genotype.

| Ancestor | Isolate | Mutation Type | Gene | Gene Function | COG Cluster |
| --- | --- | --- | --- | --- | --- |
| REL606 | 1 | - | - | - | - |
| REL606 | 2 | Missense (A43T) | *rho* | Transcription termination factor | Transcription |
| REL606 | 3 | Missense (R609S) | *topA* | DNA topoisomerase I | Replication/Repair |
| REL606 | 4 | Missense (L183Q) | *ycbC* | Hypothetical | Unknown |
| REL606 | 5 | Nonsense (E186*) | *prc* | Carboxy-terminal protease | Cell Wall/Membrane |
| REL606 | 6 | Missense (M248R) | *tldD* | Protease | Unknown |
| REL606 | 7 | Intergenic | *fabB/mnmC* | Synthase; Methyltransferase | N/A |
| REL606 | 8 | - | - | - | - |
| REL606 | 9 | - | - | - | - |
| REL606 | 10 | Missense (D99Y) | *ynfC* | Hypothetical | Unknown |
| REL606 | 11 | Nonsense (E159*) | *yfgA* | Cytoskeletal protein RodZ | Cell Wall/Membrane |
| REL606 | 12 | Missense (R334L) | *pykF* | Pyruvate kinase | Carb Metab./Trans. |
| REL606 | 13 | Deletion (157) | *infB* | Translation initiation factor IF-2 | Translation |
| REL606 | 14 | Nonsense (E159*) | *yfgA* | Cytoskeletal protein RodZ | Cell Wall/Membrane |
| REL606 | 15 | Deletion (491) | *prc* | Carboxy-terminal protease | Cell Wall/Membrane |
| REL1206 | 1 | Synonymous (V198V) | *fdhE* | Formate dehydrogenase | Post-Translation Mod. |
| REL1206 | 2 | - | - | - | - |
| REL1206 | 3 | - | - | - | - |
| REL1206 | 4 | Missense (H7Y) | *yfgA* | Cytoskeletal protein RodZ | Cell Wall/Membrane |
| REL1206 | 5 | Deletion (301) | *cls* | Cardiolipin synthetase | Lipid Metabolism |
| REL1206 | 6 | - | - | - | - |

These experiments ultimately yielded 21 beneficial mutants associated with adaptation to a minimal glucose environment. Fifteen of these mutants were derived from REL606 (8 YFP, 7 CFP) and six were derived from REL1206 (1 YFP, 5 CFP). Whole-genome sequencing revealed only a single beneficial mutation in fifteen of these mutants, while the genetic basis of adaptation was not identified in the remaining six beneficial backgrounds (Table 1; SI Text).

### Magnitude and Distribution of Direct Fitness Effects

Beneficial mutants derived from the naïve REL606 genotype produced a mean relative fitness of 1.124, which is moderately higher than what has been observed previously for beneficial mutants derived from this genotype (Ostrowski et al. 2005). However, we observed much greater fitness variance among mutants than prior studies, ranging from 1.038 to 1.291 (Figure 1). This variance was statistically significant (df=14, MS=0.0222, F=80.86, p<0.0001), with the fitness of the average mutant differing by 7.4% from the mean. No significant effect of experimental block was observed (df=3, MS=0.0002, F=0.7380, p=0.5350). In addition, because a Shapiro-Wilkes test led us to question the normality of the distribution of fitness effects (W=0.8961, p=0.0831), we performed Kruskal-Wallis tests to further evaluate effects of mutant and block on fitness. These tests confirmed a highly significant mutant effect (χ^2^=54.97, df=14, p<0.0001) and demonstrated no significant block effect (χ^2^=0.0833, df=3, p=0.9938).

**Figure 1.**
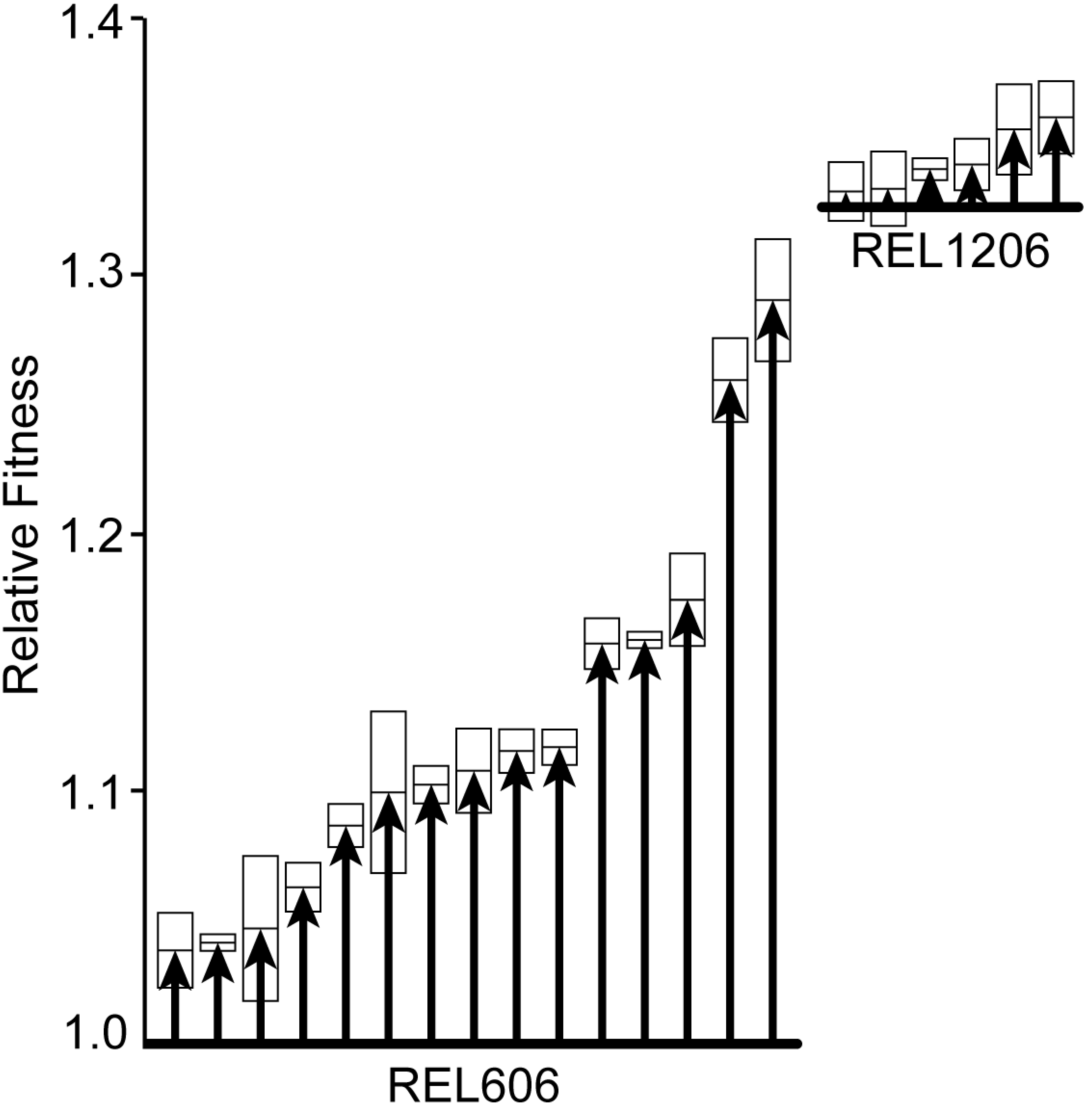
Magnitude of relative fitness increase in DM25-Glucose provided by each of the beneficial mutations isolated in this study. Each horizontal line represents the initial fitness of the founding genotype relative to the naïve REL606 ancestor across eight replicates, and each vector represents the mean fitness of each mutant relative to its founding genotype. Boxes represent 95% confidence intervals of the mean fitness across four replicates.

In contrast to the mutants from the naïve genotype, the mutants derived from glucose-adapted ancestors produced a narrow range of fitness effects, with an average of 1.014 and a range of 1.000 – 1.033 (Figure 1). Nevertheless, the variance was statistically significant (df=5, MS=0.0007, F=4.421, p=0.0113), and no significant block effect was observed (df=3, MS=0.0002, F=1.374, p=0.2889). Although the fitness values of these mutants more closely fit a normal distribution (W=0.9103, p=0.4383), we again performed Kruskal-Wallis tests on these values, which supported the significant mutant (χ^2^=12.98, df=5, p=0.0236) and non-significant block effects (χ^2^=0.7733, df=3, p=0.8558).

### Form and Distribution of Pleiotropic Fitness Effects

Pleiotropic effects of each mutant were assessed by assaying the fitness of each genotype in five alternative carbon substrates alongside glucose. Pleiotropy was both common and predominantly positive among the beneficial mutations derived from REL606 (Table 2; Figure 2). Of the fifteen REL606-derived beneficial mutants, four affected fitness in all five alternative environments, six affected fitness in four environments, four affected fitness in three environments, and one affected fitness in two environments (Table S1; Figure S4). Thus, the majority of the beneficial mutations favored during growth on glucose had significant pleiotropic effects on fitness in other resources (Table 2).

**Table 2.** Summarized results of one-sample, two-tailed t-tests in each resource for REL606 derived (top) and REL1206 derived (bottom) mutants. Mutants were designated as significantly beneficial or significantly deleterious based on a cutoff p-value of 0.05 following a Benjamini-Hochberg Correction.

| T-Test Result | Glucose | NAG | Trehalose | Galactose | Melibiose | Maltose |
| --- | --- | --- | --- | --- | --- | --- |
| Significantly Beneficial | 14 | 14 | 9 | 11 | 4 | 8 |
| Effectively Neutral | 1 | 1 | 4 | 1 | 8 | 3 |
| Significantly Deleterious | 0 | 0 | 2 | 3 | 3 | 4 |
| Significantly Beneficial | 2 | 3 | 4 | 2 | 1 | 2 |
| Effectively Neutral | 4 | 3 | 1 | 4 | 4 | 4 |
| Significantly Deleterious | 0 | 0 | 1 | 0 | 1 | 0 |

The abundant pleiotropy could be explained by systematic effects of adaptation to the laboratory environment that would produce no genotype-by-environment interactions. However, fitness variance was explained by highly significant effects of mutant (df=14, MS=0.1251, F=277.7, p<0.0001), resource (df=15, MS=0.1761, F=390.9, p<0.0001), and mutant*resource interaction (df=70, MS=0.0109, F=24.09, p<0.0001). Evidently, both mutant and resource independently affect fitness and the response of different mutants to each resource was variable. Nonetheless, the form of pleiotropy in this study was predominantly positive in all environments, and the average pleiotropic effect of REL606-derived mutants tended to be most positive in environments that shared mechanisms of inner and outer membrane transport with glucose (Table 2; Figure 2) (Travisano and Lenski 1996; Ostrowski et al. 2005). As was the case in a previous study (Ostrowski et al. 2005), we also found that when pleiotropy is positive, a significant correlation exists between the magnitude of the direct effect of a beneficial mutation and its pleiotropic effects in environments that are relatively similar to glucose (NAG, trehalose, galactose) (Table S2; Figure S5). However, we found no significant correlation between the direct fitness benefit of a mutation and its fitness in melibiose or maltose, even when antagonistic mutations were eliminated from the analysis (Table S2; Figure S5). It is worth noting that the distributions of fitness effects in melibiose and maltose were significantly non-normal (Shapiro-Wilkes test, Melibiose: W=0.8616, p=0.0254; Maltose: W=0.8632, p=0.0269). Yet Spearman’s rank correlations also demonstrated no significant relationship between the direct fitness effect in glucose and pleiotropic effects in melibiose and maltose (melibiose: df=14, p=0.4039; maltose: df=14, p=0.1779).

**Figure 2.**
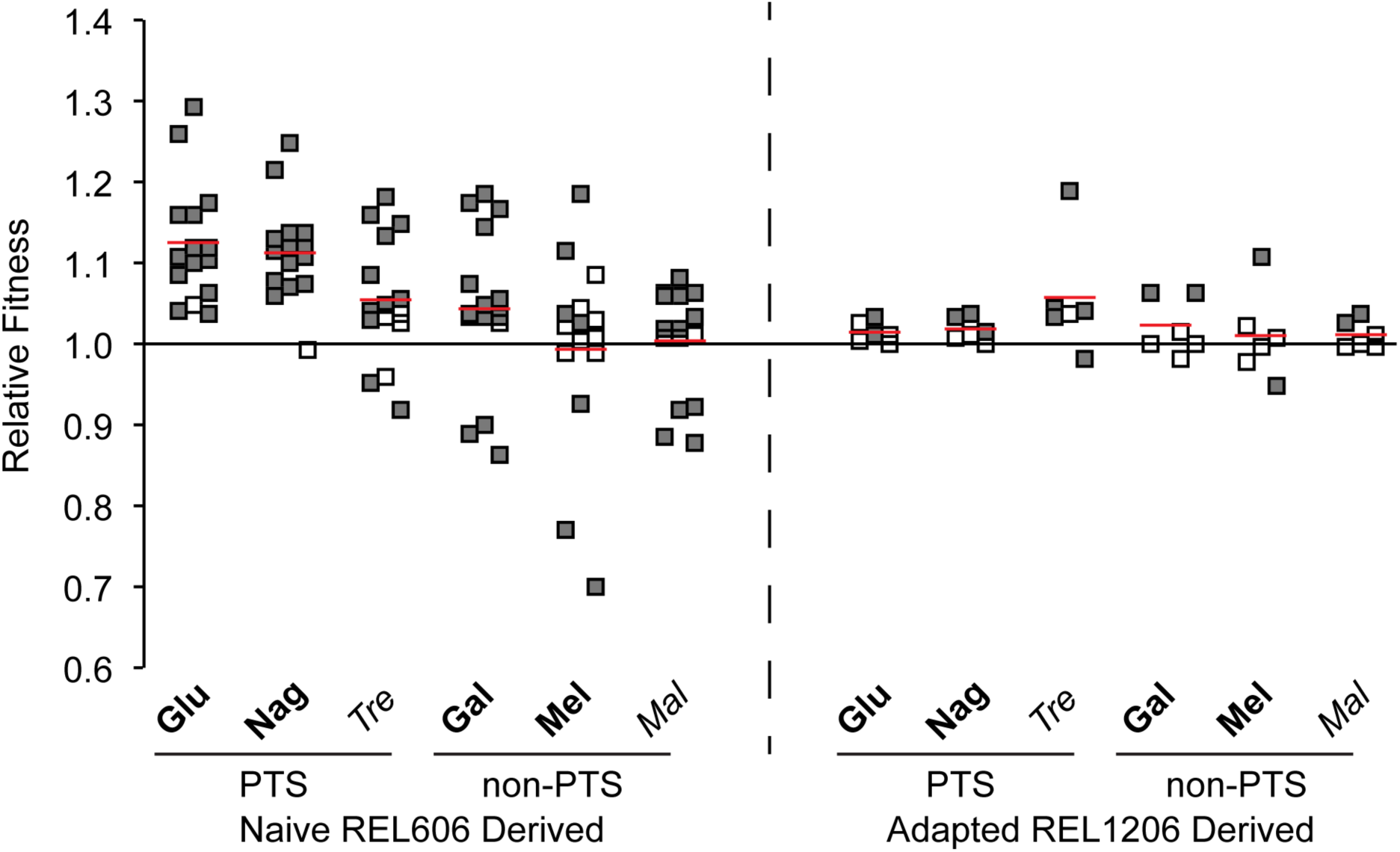
Relative fitness of each beneficial mutant against its founding genotype in DM25-Glucose (Glu) and five alternative environments: DM25-N-Acetyl-D-Glucosammine (Nag), DM25-Trehalose (Tre), DM25-Galactose (Gal), DM25-Melibiose (Mel), and DM25-Maltose (Mal). Each square represents the mean fitness of each mutant across four replicates, and each line represents the average fitness in each environment across all mutants (n = 15 for REL606; n = 6 for REL1206). Filled squares represent fitness values that are significantly different from 1, while open squares represent fitness values that are not significantly different from 1. Confidence intervals for each point are presented in Table S1. Resources are grouped by transport mechanism across the inner membrane (phosphotransferase (PTS) or non-phosphotransferase (non-PTS)) and outer membrane (OmpF (bold) or LamB (non-bold)). Each square represents the mean fitness of each mutant across four replicates, and each line represents the average fitness in each environment across all mutants (n = 15 for REL606; n = 6 for REL1206).

As was the case with direct effects of mutants derived from the glucose-adapted strain REL1206, the variance in pleiotropic effects was dramatically less than for mutants of REL606. However, pleiotropy was still common and remained predominantly positive. Of the six REL1206-derived mutants, none significantly affected fitness in all five alternative environments, but all mutants significantly affected fitness in at least one alternative environment (Table 2; Figure S4). Despite the small sample size, fitness variance was also explained by effects of mutant (df=5, MS=0.0089, F=44.94, p<0.0001), resource (df=5, MS=0.0067, F=34.03, p<0.0001), and mutant*resource interactions (df=25, MS=0.0060, F=30.62, p<0.0001). However, fitness in glucose was only positively correlated with fitness in one other environment, NAG (Table S3; Figure S6). Only fitness effects in trehalose were significantly non-normally distributed (Shapiro Wilkes: W=0.7596, p=0.0247), but Spearman’s rank correlation corroborated the non-significant relationship between the magnitude of the direct effects and pleiotropic effects in this alternative environment (df=6, p=0.6583).

While positive pleiotropy was the predominant form of pleiotropy for most mutants in this study, we observed a few cases of severely antagonistic pleiotropy. Three beneficial mutations derived from REL606 dramatically reduced fitness in all alternative environments except NAG, with declines in fitness ranging from 4 - 32% (Figure 3B). Fitness in glucose for these mutants (1.05, 1.10, and 1.12) was comparable to others producing positive pleiotropy and thus not predictive of this antagonism (Figure 1; Figure 2). Only one additional case of antagonism was seen, as one REL606 derived mutant was only deleterious in maltose (*w* = 0.92). Among mutants derived from REL1206, none were consistently antagonistic across alternative environments, but two mutants were antagonistic for fitness in a single environment (Figure 2; Table S1).

### Molecular Basis of Beneficial Mutations

As predicted by the phenotypic diversity in this study, the genetic basis of the beneficial mutations was also incredibly diverse. Importantly, no lineage contained more than one fixed mutation, which supports the inference that all phenotypes trace to a single genetic change. However, we were unable to find the genetic basis of adaptation in three of the fifteen REL606 derived beneficial mutants and three of the six REL1206 derived beneficial mutants (Table 1; Table S2).

Unlike previous studies of beneficial mutants, nearly every causative mutation disrupted a unique gene. Even at the level of functional category, mutations occurred in genes belonging to eight different COG clusters, including Carbohydrate Metabolism and Transport, Cell Wall and Membrane, Lipid Metabolism, Post-Translational Modification, Replication and Repair, Transcription, Translation, and Unknown Function. Of these COG clusters, only the Cell Wall and Membrane cluster was hit more than once and it involved all examples of gene-level parallelism (Table 1). Namely, two presumed loss of function mutations in REL606-derived mutants disrupted the *prc* gene, which encodes a carboxy-terminal protease. Further, three beneficial mutations occurred in the *yfgA* gene encoding the cytoskeletal protein RodZ, two derived from REL606 and one derived from REL1206. All other mutations affected unique genes, and only three of these genes (*pykF*, *topA,* and *infB*) have previously been identified as molecular targets of adaptation in the LTEE (Cooper et al. 2001, 2003; Woods et al. 2006; Ostrowski et al. 2008; Barrick et al. 2009; Khan et al. 2011).

### Molecular and Phenotypic Parallelism

Despite the overall lack of parallelism observed here, the two examples were especially interesting because the REL606 derived *prc* mutants exhibited the greatest fitness antagonism in alternative resources and the REL606 derived *yfgA* mutants provided the highest direct benefit in glucose. Further, a third, distinct adaptive *yfgA* mutant was isolated from the REL1206 ancestor and produced dramatically different direct and pleiotropic effects than the two REL606-derived *yfgA* mutations (Figure 3).

**Figure 3.**
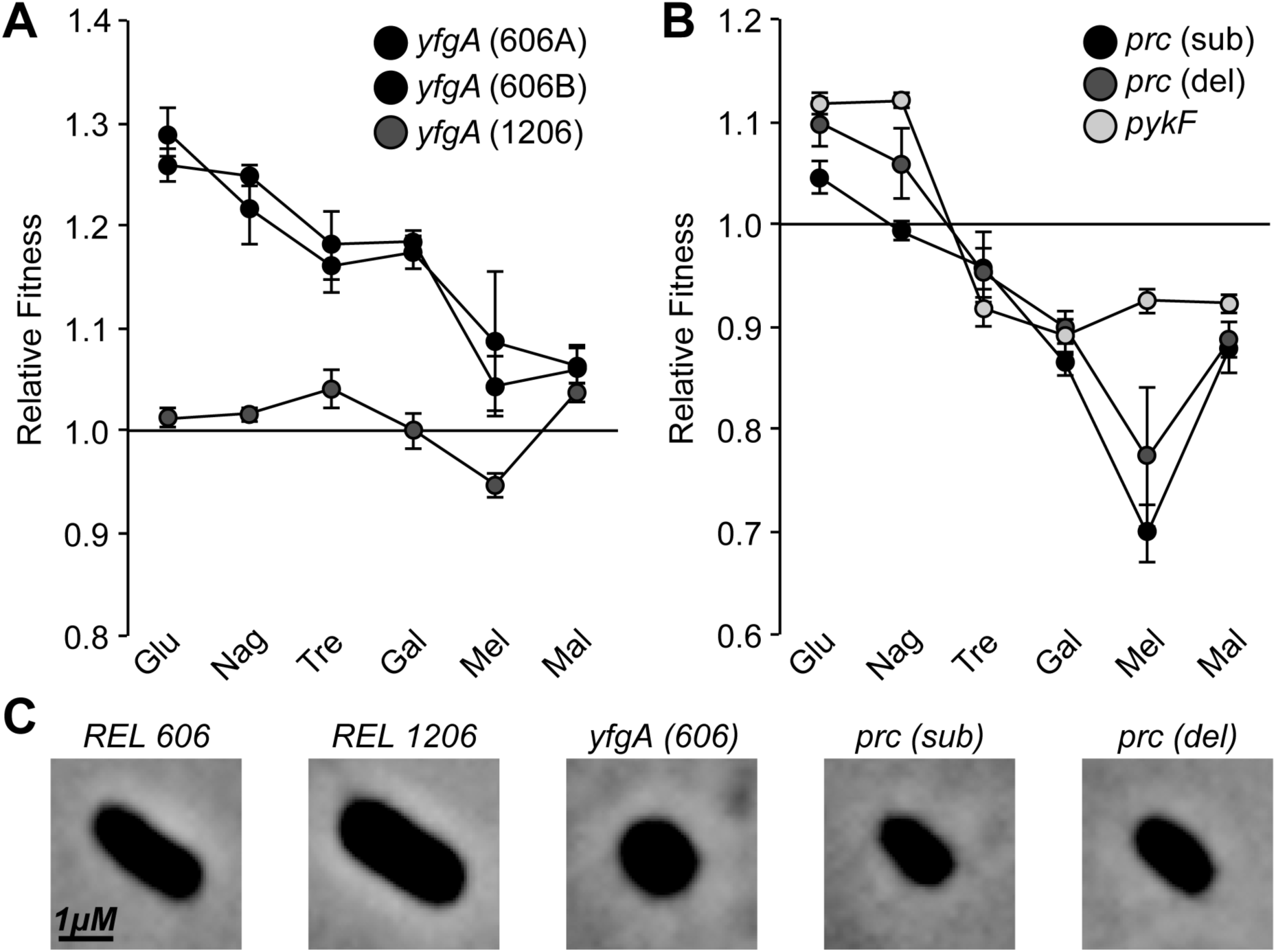
Relative fitness of parallel high benefit (A) and highly antagonistic (B) beneficial mutants in DM25-Glucose (Glu), DM25-N-Acetyl-D-Glucosammine (Nag), DM25-Trehalose (Tre), DM25-Galactose (Gal), DM25-Melibiose (Mel), and DM25-Maltose (Mal). Each point represents the average fitness across four replicates and error bars represent 95% confidence intervals of those measurements. C) Phase contrast images of both ancestral clones used in this study and the parallel cell wall biosynthesis mutants derived from REL606. All images are magnified to scale.

The two *prc* mutants derived from REL606 were distinct, with one containing a nonsense mutation (E186*) and the other containing a 1-bp deletion affecting the 491^st^ amino acid. Both of these mutations lead to a truncated product, but their fitness phenotypes are distinct, presumably because the more complete product produced by the second mutant retains partial function (Figure 3A). Interestingly, the deletion mutant predicted to produce a partial product is significantly more beneficial in glucose and reduces the pleiotropic costs of adaptation (glucose: t=4.442, df=3, p=0.0212, NAG: t=4.810, df=3, p=0.0171, and melibiose: t=11.97, df=3, p=0.0013). In contrast, the two *yfgA* mutants derived from REL606 involved genetically identical nonsense mutations (E159*). As expected, these mutants were phenotypically indistinguishable in all of the environments assayed, produced the greatest fitness increase in glucose, and were positively pleiotropic (Figure 3B). This *yfgA* mutation demonstrates that large gains in fitness can be achieved with few pleiotropic costs (Figure 2). The third *yfgA* mutation (H7Y) that occurred in the REL1206 background was distinct, producing a missense mutation rather than nonsense. Interestingly, both *prc* and *yfgA* mutations produced truncated, smaller cells (Figure 3C) despite the dramatic differences in their fitness effects.

Lastly, three mutants evolved from REL606 acquired mutations in *pykF*, *topA*, and *infB*, genes that have been identified as targets of selection in the LTEE. The nonsense mutation in the *pykF* gene (R334L) was the third highly antagonistically pleiotropic mutant and reduced fitness in all alternative environments except NAG (Figure 3B). The nonsense mutation in *topA* (R609S) was generally positively pleiotropic as it increased fitness in all environments except melibiose (Table S1).

Lastly, the 1-bp deletion in the *infB* allele resulted in a dramatic increase in direct fitness of 17.44%. This mutant was positively pleiotropic in NAG, trehalose, and galactose, but was neutral in melibiose and deleterious in maltose (Table S1).

## DISCUSSION

Despite their importance in countless biological processes, studies addressing the nature and pleiotropic effects of beneficial mutations remain limited (Elena and Lenski 2003; Orr 2003; Kassen and Bataillon 2006; Eyre-Walker and Keightley 2007; Bataillon et al. 2011; Good et al. 2012; Rice et al. 2015). Here we built upon established methods for the detection and analysis of beneficial mutations in different population genetic environments by using current technologies for cell counting and sequencing. This approach enabled a diverse collection of beneficial mutants that includes some of the variation that may be typically lost due to random sampling and clonal interference in large populations (Gerrish and Lenski 1998; Ostrowski et al. 2005; Levy et al. 2015). We determined the direct and pleiotropic fitness effects of these mutants in six environments and identified the genetic basis of adaptation using whole-genome sequencing.

In an attempt to broaden the sampled diversity of beneficial mutations, we decreased the effective population size of our short-term selection experiments, isolated beneficial mutations from both a naïve and glucose-adapted ancestor, and screened cells from both the winning and losing populations to produce our final collection of 21 single-step beneficial mutations (Gerrish and Lenski 1998; Wahl and Gerrish 2001; de Visser and Rozen 2005). As expected, this resulted in diverse phenotypic responses to selection in DM25-glucose, with an average direct fitness effect of 1.124 for mutants derived from the naïve ancestor (REL606) and an average direct fitness effect of 1.014 for mutants derived from the glucose-adapted ancestor (REL1206). Individual mutants of REL606 differed from this fitness mean by 7.4%, while mutants of REL1206 differed from the overall mean by 1.2%. In contrast, a previous collection of 27 beneficial mutants collected from REL606 at higher N_E_ exhibited a fitness deviation of only 0.026%, despite having a similar overall mean fitness to our mutants (1.096) (Ostrowski et al. 2005). Although there is no comparative dataset for our mutants derived from REL1206, theory and experiments support the decreased magnitude of fitness benefits of mutations in adapted genotypes owing to diminishing returns epistasis (Khan et al. 2011; Good et al. 2012; Kryazhimskiy et al. 2014; Nahum et al. 2015).

Although the predominant form of pleiotropy observed in this study was positive, pleiotropic effects were also diverse. Most mutants exhibited universally positive pleiotropy in the five alternative environments, but others exhibited broadly antagonistic pleiotropy. While many evolutionary models have assumed that pleiotropy is mostly antagonistic (Fisher 1930; Lande 1983; Orr and Coyne 1992; Otto 2004), positive pleiotropy seems to be the rule rather than the exception when beneficial mutants are assayed on alternative carbon substrates (Ostrowski et al. 2005; Lee et al. 2009). Prior studies have also identified that adaptation can be resource-specific and can influence the magnitude and direction of pleiotropy in this system. For example, resources that share similar mechanisms of inner and outer membrane transport with glucose have been shown to permit similar fitness levels, while resources with dissimilar transport mechanisms reveal neutral or antagonistic effects (Travisano et al. 1995b; Travisano and Lenski 1996; Ostrowski et al. 2005). Adaptation appears to be resource-specific among the REL606-derived mutations studied here (Figure 2), but we lack enough statistical power to identify any correlation with resource transport for mutants derived from REL1206. However, some REL1206 mutants achieved fitness in foreign environments greater than in glucose, despite little similarity in the resource transport mechanism (Table S1). This result may indicate that the targets of adaptation have changed over the course of 2,000 generations of prior adaptation to glucose. Furthermore, while antagonistic pleiotropy was relatively rare (Ostrowski et al. 2005; Lee et al. 2009), it was almost exclusively limited to a subset of three particular mutants (Figure 3B), suggesting that different paths of adaptation yielding similar fitness benefits in the selective environment may produce vastly different evolutionary trade-offs (Rodriguez-Verdugo et al. 2014).

The genetic basis of the beneficial mutations collected in this study was also diverse and does not support the oligogenic model of adaptation discussed in previous studies (Wood et al. 2005; Ostrowski et al. 2008). Among 15 beneficial mutants tracing to a single, known mutation, only two cases of parallelism at the gene level were observed (Table 1). Moreover, we isolated mutations in only three genes that were previously identified as targets of adaptation in the LTEE (*pykF*, *topA*, and *infB*), despite the analysis of several thousand generations of whole-genome sequencing data from this project (Cooper et al. 2001, 2003; Woods et al. 2006; Ostrowski et al. 2008; Barrick et al. 2009; Khan et al. 2011). The remaining twelve mutants represent novel selective targets for adaptation in an environment that closely resembles that of the LTEE. These novel mutations occurred in genes belonging to six different COG clusters, and include missense, nonsense, synonymous, and indel mutations (Table 1). Collectively, this study suggests that that the number of mutations producing detectable benefit under this selective regime may be greater than previously expected, and this increase may be a product of sampling from populations evolved at lower population size and N_E_. In addition, subtle differences in our model such as the reduced culture volume and altered oxygenation from growing in test tubes may explain the novel mutational targets. It seems likely that the five mutants with relative fitness improvements greater than 10% (with mean fitness ranging from 1.14 to 1.29) could represent new targets of selection in our model as compared to the LTEE, in which single mutations producing benefits greater than 10% were rarely observed (Lenski et al. 1991; Lenski and Travisano 1994; Barrick et al. 2009; Wiser et al. 2013).

A few key generalizations can be made from the adaptive genotypes and their fitness effects. First, the direct effects of beneficial mutations derived from naïve genotypes on glucose are substantially higher than those derived from glucose-adapted genotypes (Figure 1). Second, the pleiotropic effects of beneficial mutations on alternative carbon substrates are generally positive from both ancestors, but pleiotropic effects have a greater magnitude when derived from naïve genotypes (Figure 2). Lastly, mutations in a variety of genes can evidently produce high benefit in this selective regime, but this number is decreased in adapted genotypes (Table 1; Figure S2; Figure S3). These findings broadly agree with Fisher’s geometric model of adaptation, which assumes that pleiotropy is universal and hence large-effect mutations would be more pleiotropic than small-effect mutations (Blanquart et al. 2014; Matuszewski et al. 2014).

Exceptions to these generalizations are equally important as they can have dramatic consequences on evolving populations. The two genes in which more than one beneficial mutation was detected produced effects that disagree with these patterns. In one case, mutants of *yfgA* produced large direct benefits without any pleiotropic costs, and in the other case, mutants of *prc* produced highly antagonistic effects with only moderate direct benefits. Three different beneficial mutants affected *yfgA*, which encodes the cytoskeletal protein RodZ, a key component of cell wall biosynthesis. Both REL606-derived *yfgA* mutations were genetically identical nonsense mutations (E159*) that produced the greatest direct fitness benefit (1.29 and 1.26) of all mutants isolated in this study and displayed positive or neutral pleiotropic effects in all alternative environments (Figure 3A). A third *yfgA* missense mutation (H7Y) was isolated from REL1206 that produced a fitness gain of only 1.012 and with limited pleiotropic effects (Figure 3A). Individual mutations producing a selective benefit of greater than 25% are expected to be extremely rare, as illustrated by the fact that single *yfgA* mutants in REL606 nearly reached the relative fitness of REL1206, which harbors at least five beneficial mutations over the course of 2000 generations of adaptation to glucose (Figure 1; SI Text). This result implies that single beneficial mutations can produce large fitness gains with limited pleiotropic costs. The lesser advantage of the *yfgA* mutant of REL1206 in both selective and alternative environments could be explained either by its less severe missense mutation or by diminishing returns epistasis (Khan et al. 2011; Kryazhimskiy et al. 2014). However, it is also noteworthy that this *yfgA* missense mutation was deleterious in melibiose (Figure 3; Table S1), suggesting that both the magnitude and direction of pleiotropy can vary as a result of genetic background (Flynn et al. 2013). The two *prc* mutants, encoding a periplasmic carboxyl terminal protease, involve a nonsense substitution (E186*) and a 1bp deletion (Table 1). Along with a missense mutation in *pykF* (R334L), which encodes a pyruvate kinase, these three mutants account for eleven of the twelve cases of significantly antagonistic pleiotropy observed in REL606-derived mutants. Each mutant was antagonistically pleiotropic in all three non-PTS resources, with relative fitness ranging from 0.70 to 0.92, and the *prc* nonsense and *pykF* mutants were also significantly deleterious in trehalose (Figure 3B; Table S1).

Despite the opposing pleiotropic effects of mutations in *yfgA* and *prc*, both genes are involved in cell wall biosynthesis and phase-contrast microscopy reveals similar impacts on cell morphology (Figure 3C). The *yfgA* gene encodes a cytoskeletal protein that helps to maintain the rigid rod morphology of *E. coli* cells by anchoring to the cytoplasmic membrane and interacting with penicillin binding proteins (PBPs) to help synthesize the peptidoglycan layer of the cell wall (Jeong et al. 2009; Philippe et al. 2009). The *yfgA* product interacts with MreB to form the actin cytoskeleton and constrain peptidoglycan synthesis to the periplasm (Shiomi et al. 2008; van den Ent et al. 2010). Interestingly, *mreB* mutations can confer a similar fitness benefit via cell size changes in *E. coli* (Monds et al. 2014). The *prc* carboxyl terminal protease activates PBPs to synthesize the peptidoglycan layer (Hara et al. 1991; Tadokoro et al. 2004). Loss of function mutations in *yfgA, mreB*, and *prc* would therefore be predicted to synthesize a more limited peptidoglycan layer and thus produce more spherical cells. Microscopy of these *yfgA* and *prc* mutants supported this prediction (Figure 3C). These results add to numerous reports of beneficial changes in cell morphology in the LTEE, where loss in rod-shape morphology and cell size increases appear to scale linearly with fitness (Philippe et al. 2009; Monds et al. 2014). Although these changes in cell morphology are expected to be highly pleiotropic via their affects on nutrient uptake, cell survival, and growth, mutations in the *prc* gene may be more predisposed to antagonistic pleiotropy because of its involvement in the activation of other proteins along with PBPs (Hara et al. 1991; Tadokoro et al. 2004). In sum, although the amount of genetic parallelism observed in this study was limited, the few examples we did observe suggest that strong selection on a particular target may nevertheless produce highly divergent evolution in niche breadth, with some mutations resulting in pleiotropic gains in fitness, and others resulting in pleiotropic loses.

Despite growing evidence of the universality and diversity of pleiotropy, which is the root cause of tradeoffs produced by selection, we still know relatively little about the commonality and the form of pleiotropic effects of beneficial mutations, even in well-characterized systems (Cooper and Lenski 2000; Cooper et al. 2001; Dudley et al. 2005; Ostrowski et al. 2005; Kassen and Bataillon 2006). We have shown that a diverse collection of beneficial mutations can generate highly divergent pleiotropic outcomes. On average, the magnitude of pleiotropic effects scaled with the magnitude of the fitness increase in the selected environment, and hence mutations affecting the less adapted ancestor were more pleiotropic, and mutations affecting the more adapted ancestor were less pleiotropic. These overall patterns tend to support Fisher’s geometric model of adaptation, in which pleiotropy is universal (Orr 2006; Blanquart et al. 2014; Matuszewski et al. 2014). However, the form of pleiotropy was generally positive, which disagrees with many tradeoff models predicated by the geometric model (Orr 2000; Otto 2004; Lourenco et al. 2011). Nevertheless, some mutants that provided a direct benefit on glucose were highly antagonistic on other carbon sources. Ultimately, these findings suggest that many beneficial variants are accessible at different points on the fitness landscape, and the success of a particular beneficial mutation can either expand the fundamental niche through positive pleiotropy, or lead to specialization via loss of fitness on alternative carbon substrates. However, while our mutants were collected in a simple environment with a single carbon source and constant temperature, nature surely presents greater environmental heterogeneity. Thus, which of these evolutionary paths is followed, and the eventual shape of the niche, may depend on the influence of environmental fluctuations that expose previously hidden pleiotropic effects that favor or disfavor contending mutations.

## Acknowledgements

We thank all members of the Cooper lab for helpful discussion and Bob Mooney for microscopic support. This work was supported by the National Science Foundation Career Award (DEB-0845851 to VSC).

## Literature Cited

Andrews, S. 2010. FastQC: A quality control tool for high throughput sequence data.

Barrick, J. E., and R. E. Lenski. 2009. Genome-wide mutational diversity in an evolving population of *Escherichia coli*. Cold Spring Harb. Symp. Quant. Biol. 74:119–129.

Barrick, J. E., D. S. Yu, S. H. Yoon, H. Jeong, T. K. Oh, D. Schneider, R. E. Lenski, and J. F. Kim. 2009. Genome evolution and adaptation in a long-term experiment with *Escherichia coli*. Nature 461:1243–1247.

Bataillon, T., T. Zhang, and R. Kassen. 2011. Cost of adaptation and fitness effects of beneficial mutations in *Pseudomonas fluorescens*. Genetics 189:939–949.

Baym, M., S. Kryazhimskiy, T. D. Lieberman, H. Chung, M. M. Desai, and R. Kishony. 2015. Inexpensive multiplexed library preparation for megabase-sized genomes. PLoS ONE 10:e0128036.

Benjamini, Y., and Y. Hochberg. 1995. Controlling the false discovery rate: a practical and powerful approach to multiple testing. J. R. Stat. Soc. 57:289–300.

Blanquart, F., G. Achaz, T. Bataillon, and O. Tenaillon. 2014. Properties of selected mutations and genotypic landscapes under Fisher’s geometric model. 68:3537–3554.

Bolger, A. M., M. Lohse, and B. Usadel. 2014. Trimmomatic: A flexible trimmer for Illumina sequence data. Bioinformatics 30:2114–2120.

Chevin, L.-M. 2011. On measuring selection in experimental evolution. Biol. Lett. 7:210– 213.

Cooper, T. F., D. E. Rozen, and R. E. Lenski. 2003. Parallel changes in qene expression after 20,000 generations of evolution in *Escherichia coli*. Proc. Natl. Acad. Sci. U. S. A. 100:1072–1077.

Cooper, V. S. 2002. Long-term experimental evolution in *Escherichia coli*. X. Quantifying the fundamental and realized niche. BMC Evol. Biol. 2:12.

Cooper, V. S. 2014. The origins of specialization: Insights from bacteria held 25 years in captivity. PLoS Biol. 12:e1001790.

Cooper, V. S., and R. E. Lenski. 2000. The population genetics of ecological specialization in evolving *Escherichia coli* populations. Nature 407:736–739.

Cooper, V. S., D. Schneider, M. Blot, and R. E. Lenski. 2001. Mechanisms causing rapid and parallel losses of ribose catabolism in evolving populations of *Escherichia coli* B. J. Bacteriol. 183:2834–2841.

Datsenko, K. A., and B. L. Wanner. 2000. One-step inactivation of chromosomal genes in *Escherichia coli* K-12 using PCR products. Proc. Natl. Acad. Sci. U. S. A. 97:6640– 6645.

De Visser, J. A. G. M., and D. E. Rozen. 2006. Clonal interference and the periodic selection of new beneficial mutations in Escherichia coli. Genetics 172:2093–2100.

De Visser, J. A. G. M., and D. E. Rozen. 2005. Limits to adaptation in asexual populations. J. Evol. Biol. 18:779–788.

Deatherage, D. E., and J. E. Barrick. 2014. Identification of mutations in laboratory-evolved microbes from next-generation sequencing data using breseq. Methods Mol. Biol. 1151:165–188.

Dudley, A. M., D. M. Janse, A. Tanay, R. Shamir, and G. M. Church. 2005. A global view of pleiotropy and phenotypically derived gene function in yeast. Mol. Syst. Biol. 1:1–11.

Elena, S. F., and R. E. Lenski. 2003. Evolution experiments with microorganisms: The dynamics and genetic bases of adaptation. Nat. Rev. Genet. 4:457–469.

Eyre-Walker, A., and P. D. Keightley. 2007. The distribution of fitness effects of new mutations. Nat. Rev. Genet. 8:610–8.

Feist, A. M., C. S. Henry, J. L. Reed, M. Krummenacker, A. R. Joyce, P. D. Karp, L. J. Broadbelt, V. Hatzimanikatis, and B. Ø. Palsson. 2007. A genome-scale metabolic reconstruction for *Escherichia coli* K-12 MG1655 that accounts for 1260 ORFs and thermodynamic information. Mol. Syst. Biol. 3:121.

Fisher, R. A. 1930. The genetical theory of natural selection. Second Edi. Oxford University Press, Oxford.

Flynn, K. M., T. F. Cooper, F. B. G. Moore, and V. S. Cooper. 2013. The environment affects epistatic interactions to alter the topology of an empirical fitness landscape. PLoS Genet. 9:e1003426.

Gallet, R., T. F. Cooper, S. F. Elena, and T. Lenormand. 2012. Measuring selection coefficients below 10(-3): Method, questions, and prospects. Genetics 190:175–186.

Gerrish, P. J., and R. E. Lenski. 1998. The fate of competing beneficial mutations in an asexual population. Genetica 102/103:127–144.

Good, B. H., I. M. Rouzine, D. J. Balick, O. Hallatschek, and M. M. Desai. 2012. Distribution of fixed beneficial mutations and the rate of adaptation in asexual populations. Proc. Natl. Acad. Sci. 109:4950–4955.

Hara, H., Y. Yamamoto, A. Higashitani, H. Suzuki, and Y. Nishimura. 1991. Cloning, mapping, and characterization of the *Escherichia coli* prc gene, which is involved in C-terminal processing of penicillin-binding protein 3. J. Bacteriol. 173:4799–4813.

Hietpas, R. T., C. Bank, J. D. Jensen, and D. N. A. Bolon. 2013. Shifting fitness landscapes in response to altered environments. Evolution (N. Y). 67:3512–3522.

Holt, R. D. 1996. Demographic constraints in evolution: Towards unifying the evolutionary theories of senescence and niche conservatism. Evol. Ecol. 10:1–11.

Jasmin, J. N., and C. Zeyl. 2013. Evolution of pleiotropic costs in experimental populations. J. Evol. Biol. 26:1363–1369.

Jeong, H., V. Barbe, C. H. Lee, D. Vallenet, D. S. Yu, S. H. Choi, A. Couloux, S. W. Lee, S. H. Yoon, L. Cattolico, C. G. Hur, H. S. Park, B. Ségurens, S. C. Kim, T. K. Oh, R. E. Lenski, F. W. Studier, P. Daegelen, and J. F. Kim. 2009. Genome Sequences of Escherichia coli B strains REL606 and BL21(DE3). J. Mol. Biol. 394:644–652.

Kassen, R., and T. Bataillon. 2006. Distribution of fitness effects among beneficial mutations before selection in experimental populations of bacteria. Nat. Genet. 38:484– 488.

Khan, A. I., D. M. Dinh, D. Schneider, R. E. Lenski, and T. F. Cooper. 2011. Negative epistasis between beneficial mutations in an evolving bacterial population. Science 332:1193–1196.

Kryazhimskiy, S., D. P. Rice, E. R. Jerison, and M. M. Desai. 2014. Global epistasis makes adaptation predictable despite sequence-level stochasticity. Science 344:1519– 1522.

Lande, R. 1983. The response to selection on major and minor mutations affecting a metrical trait. Heredity (Edinb). 50:47–65.

Le Gac, M., T. F. Cooper, S. Cruveiller, C. Médigue, and D. Schneider. 2013. Evolutionary history and genetic parallelism affect correlated responses to evolution. Mol. Ecol. 22:3292–3303.

Lee, M. C., H. H. Chou, and C. J. Marx. 2009. Asymmetric, bimodal trade-offs during adaptation of methylobacterium to distinct growth substrates. Evolution (N. Y). 63:2816– 2830.

Leiby, N., and C. J. Marx. 2014. Metabolic erosion primarily through mutation accumulation, and not tradeoffs, drives limited evolution of substrate specificity in *Escherichia coli*. PLoS Biol. 12:e1001789.

Lenski, R. E., M. R. Rose, S. C. Simpson, and S. C. Tadler. 1991. Long-term experimental evolution in *Escherichia coli* .1. Adaptation and divergence during 2,000 generations. Am. Nat. 138:1315–1341.

Lenski, R. E., and M. Travisano. 1994. Dynamics of adaptation and diversification - a 10,000-generation experiment with bacterial-populations. Proc. Natl. Acad. Sci. U. S. A. 91:6808–6814.

Levy, S. F., J. R. Blundell, S. Venkataram, D. a. Petrov, D. S. Fisher, and G. Sherlock. 2015. Quantitative evolutionary dynamics using high-resolution lineage tracking. Nature 519:181–186.

Lourenco, J., N. Galtier, and S. Glemin. 2011. Complexity, pleiotropy, and the fitness effect of mutations. Evolution (N. Y). 65:1559–1571.

Martin, G., and T. Lenormand. 2008. The distribution of beneficial and fixed mutation fitness effects close to an optimum. Genetics 179:907–916.

Matuszewski, S., J. Hermisson, and M. Kopp. 2014. Fisher’s geometric model with a moving optimum.

Monds, R. D., T. K. Lee, T. F. Cooper, K. C. Huang, R. D. Monds, T. K. Lee, A. Colavin, T. Ursell, S. Quan, and T. F. Cooper. 2014. Systematic perturbation of cytoskeletal function reveals a linear scaling relationship between cell geometry and fitness. CellReports 9:1528–1537.

Nahum, J. R., P. Godfrey-Smith, B. N. Harding, J. H. Marcus, J. Carlson-Stevermer, and B. Kerr. 2015. A tortoise–hare pattern seen in adapting structured and unstructured populations suggests a rugged fitness landscape in bacteria. Proc. Natl. Acad. Sci. 112:7530–7535.

Orr, H. A. 2000. Adaptation and the cost of complexity. Evolution (N. Y). 54:13–20.

Orr, H. A. 2003. The distribution of fitness effects among beneficial mutations. Genetics 163:1519–1526.

Orr, H. A. 2006. The distribution of fitness effects among beneficial mutations in Fisher’s geometric model of adaptation. J. Theor. Biol. 238:279–285.

Orr, H. A., and J. A. Coyne. 1992. The genetics of adaptation: a reassessment. Am. Nat. 140:725–742.

Ostrowski, E. A., D. E. Rozen, and R. E. Lenski. 2005. Pleiotropic effects of beneficial mutations in *Escherichia coli*. Evolution (N. Y). 59:2343–2352.

Ostrowski, E. A., R. J. Woods, and R. E. Lenski. 2008. The genetic basis of parallel and divergent phenotypic responses in evolving populations of *Escherichia coli*. Proc. Biol. Sci. 275:277–284.

Otto, S. P. 2004. Two steps forward, one step back: the pleiotropic effects of favoured alleles. Proc. R. Soc. London Ser. B-Biological Sci. 271:705–714.

Perfeito, L., a. Sousa, T. Bataillon, and I. Gordo. 2014. Rates of fitness decline and rebound suggest pervasive epistasis. Evolution (N. Y). 68:150–162.

Philippe, N., L. Pelosi, R. E. Lenski, and D. Schneider. 2009. Evolution of penicillin-binding protein 2 concentration and cell shape during a long-term experiment with *Escherichia coli*. J. Bacteriol. 191:909–921.

R Development Core Team. 2013. R: A Language and Environment for Statistical Computing.

Remold, S. K., and R. E. Lenski. 2001. Contribution of individual random mutations to genotype-by-environment interactions in *Escherichia coli*. Proc. Natl. Acad. Sci. U. S. A. 98:11388–11393.

Rice, D. P., B. H. Good, and M. M. Desai. 2015. The evolutionarily stable distribution of fitness effects. Genetics 200:321–329.

Rodriguez-Verdugo, A., D. Carrillo-Cisneros, A. Gonzalez-Gonzalez, B. S. Gaut, and A. F. Bennett. 2014. Different tradeoffs result from alternate genetic adaptations to a common environment. Proc. Natl. Acad. Sci. 111:12121–12126.

Rozen, D. E., J. A. G. M. de Visser, and P. J. Gerrish. 2002. Fitness effects of fixed beneficial mutations in microbial populations. Curr. Biol. 12:1040–1045.

Shiomi, D., M. Sakai, and H. Niki. 2008. Determination of bacterial rod shape by a novel cytoskeletal membrane protein. EMBO J. 27:3081–3091.

Tadokoro, A., H. Hayashi, T. Kishimoto, Y. Makino, S. Fujisaki, and Y. Nishimura. 2004. Interaction of the *Escherichia coli* lipoprotein NlpI with periplasmic Prc (Tsp) protease. J. Biochem. 135:185–191.

Tenaillon, O., A. Rodriguez-Verdugo, R. L. Gaut, P. McDonald, A. F. Bennett, A. D. Long, and B. S. Gaut. 2012. The molecular diversity of adaptive convergence. Science 335:457–461.

Travisano, M., and R. E. Lenski. 1996. Long-term experimental evolution in Escherichia coli. 4. Targets of selection and the specificity of adaptation. Genetics 143:15–26.

Travisano, M., J. A. Mongold, A. F. Bennett, and R. E. Lenski. 1995a. Experimental tests of the roles of adaptation, chance, and history in evolution. Science 267:87–90.

Travisano, M., F. Vasi, and R. E. Lenski. 1995b. Long-term experimental evolution in *Escherichia coli* .3. Variation among replicate populations in correlated responses to novel environments. Evolution (N. Y). 49:189–200.

Van den Ent, F., C. M. Johnson, L. Persons, P. de Boer, and J. Löwe. 2010. Bacterial actin MreB assembles in complex with cell shape protein RodZ. EMBO J. 29:1081– 1090.

Wahl, L. M., and P. J. Gerrish. 2001. The probability that beneficial mutations are lost in populations with periodic bottlenecks. Evolution (N. Y). 55:2606–2610.

Wiser, M. J., N. Ribeck, and R. E. Lenski. 2013. Long-term dynamics of adaptation in asexual populations. Science 342:1364–1367.

Wood, T. E., J. M. Burke, and L. H. Rieseberg. 2005. Parallel genotypic adaptation: When evolution repeats itself. Genetica 123:157–170.

Woods, R., D. Schneider, C. L. Winkworth, M. A. Riley, and R. E. Lenski. 2006. Tests of parallel molecular evolution in a long-term experiment with *Escherichia coli*. Proc. Natl. Acad. Sci. U. S. A. 103:9107–9112.

Treutlein, B. et al. Reconstructing lineage hierarchies of the distal lung epithelium using single-cell RNA-seq. Nature 509, 371–5. issn: 1476-4687 (May 2014).

Buettner, F. et al. Computational analysis of cell-to-cell heterogeneity in single-cell RNA-sequencing data reveals hidden subpopulations of cells. Nature Biotechnology. issn: 1087-0156. doi:10.1038/nbt.3102. http://www.nature.com/doifinder/10.1038/nbt. 3102 (Jan. 2015).

Moignard, V. et al. Decoding the regulatory network of early blood development from single-cell gene expression measurements. Nature Biotechnology 33. issn: 1087-0156. doi:10.http://dx.doi.org/10.1038/nbt.3154”1038/nbt.3154. http://www.nature.com/doifinder/10.1038/nbt.3154 (Feb. 2015).

Macaulay, I. C. & Voet, T. Single cell genomics: advances and future perspectives. PLoS genetics 10, e1004126. issn: 1553-7404 (Jan. 2014).

Stegle, O., Teichmann, S. a. & Marioni, J. C. Computational and analytical challenges in single-cell transcriptomics. Nature Reviews Genetics 16, 133–145. issn: 1471-0056 (Jan. 2015).

Trapnell, C. et al. The dynamics and regulators of cell fate decisions are revealed by pseudotemporal ordering of single cells. Nature biotechnology 32, 381–6. issn: 1546-1696 (Apr. 2014).

Hyvärinen, A. Fast and robust fixed-point algorithms for independent component analysis. IEEE transactions on neural networks / a publication of the IEEE Neural Networks Council 10, 626–34. issn: 1045-9227 (Jan. 1999).

Pandey, S., Billor, N. & Turkmen, A. The Effect of Outliers in Independent Component Analysis. American Journal of Mathematical and Management Sciences 28, 399–418. issn: 0196-6324 (Feb. 2008).

Kharchenko, P. V., Silberstein, L. & Scadden, D. T. Bayesian approach to single-cell differential expression analysis. Nature methods 11, 740–2. issn: 1548-7105 (July 2014).

Bendall, S. C. et al. Single-cell trajectory detection uncovers progression and regulatory coordination in human B cell development. Cell 157, 714–25. issn: 1097-4172 (Apr. 2014).

Shin, J. et al. Single-Cell RNA-Seq with Waterfall Reveals Molecular Cascades underlying Adult Neurogenesis. English. Cell Stem Cell 17, 360–372. issn: 1934-5909 (Aug. 2015).

Haghverdi, L., Buettner, F. & Theis, F. J. Diffusion maps for high-dimensional single-cell analysis of differentiation data. Bioinformatics, 1–10. issn: 1367-4803 (2015).

Belkin, M. & Niyogi, P. Laplacian Eigenmaps for Dimensionality Reduction and Data. 1396, 1373–1396 (2003).

Luxburg, U. A tutorial on spectral clustering. Statistics and Computing 17, 395–416. issn: 0960-3174 (Aug. 2007).

Hastie, T. & Stuetzle, W. Principal Curves. en. http://www.tandfonline.com/doi/abs/10.1080/01621459.1989.10478797 (Mar. 2012).

Brennecke, P. et al. Accounting for technical noise in single-cell RNA-seq experiments. Nature methods 10, 1093–5. issn: 1548-7105 (Nov. 2013).

Trapnell, C. et al. Transcript assembly and quantification by RNA-Seq reveals unannotated transcripts and isoform switching during cell differentiation. Nature biotechnology 28, 511–5. issn: 1546-1696 (May 2010).

Young, M. D., Wakefield, M. J., Smyth, G. K. & Oshlack, A. Gene ontology analysis for RNA-seq: accounting for selection bias. Genome biology 11, R14. issn: 1465-6914 (Jan. 2010).

Julia, M., Telenti, A. & Rausell, A. Sincell: an R/Bioconductor package for statistical as- sessment of cell-state hierarchies from single-cell RNA-seq. Bioinformatics, btv368–. issn: 1367-4803 (June 2015).

